# Neurons from individual early Alzheimer’s disease patients reflect their clinical vulnerability

**DOI:** 10.1101/2021.11.09.467891

**Authors:** Bryan Ng, Helen A. Rowland, Tina Wei, Kanisa Arunasalam, Emma Mee Hayes, Ivan Koychev, Anne Hedegaard, Elena M. Ribe, Dennis Chan, Tharani Chessell, Dominic ffytche, Roger N. Gunn, Ece Kocagoncu, Jennifer Lawson, Paresh Malhotra, Basil H. Ridha, James B. Rowe, Alan J. Thomas, Giovanna Zamboni, Noel J. Buckley, Zameel M. Cader, Simon Lovestone, Richard Wade-Martins

## Abstract

Establishing preclinical models of Alzheimer’s disease that predict clinical outcomes remains a critically important, yet to date not fully realised, goal. Models derived from human cells offer considerable advantages over non-human models, including the potential to reflect some of the inter-individual differences that are apparent in patients. Here we report an approach using induced pluripotent stem cell-derived cortical neurons from people with early symptomatic Alzheimer’s disease where we sought a match between individual disease characteristics in cells with analogous characteristics in the people from whom they were derived. We show that the response to amyloid-β burden *in life,* as measured by cognitive decline and brain activity levels, varies between individuals and this vulnerability rating correlates with the individual cellular vulnerability to extrinsic amyloid-β *in vitro* as measured by synapse loss and function. Our findings indicate that patient induced pluripotent stem cell-derived cortical neurons not only present key aspects of Alzheimer’s disease pathology, but also reflect key aspects of the clinical phenotypes of the same patients. Cellular models that reflect an individual’s in-life clinical vulnerability thus represent a tractable method of Alzheimer’s disease modelling using clinical data in combination with cellular phenotypes.

## Introduction

Alzheimer’s disease is the most common age-related neurodegenerative disease and cause of dementia, estimated to affect close to 50 million people in 2015 worldwide with cases predicted to almost double every 20 years^1^. Autosomal dominant mutations in the Amyloid Precursor Protein gene (*APP*) or genes encoding the APP proteolytic enzymes Presenilins 1 and 2 (*PSEN1, PSEN2*) are causative of early onset familial Alzheimer’s disease. Largely based on insights from familial Alzheimer’s disease, amyloid-β (Aβ) generation, metabolism or clearance is thought to underlie the pathogenesis of late onset forms of sporadic Alzheimer’s disease. However, it is also apparent that whilst amyloid-related features predict clinical outcomes, this relationship shows very considerable inter-individual variation^2^. Some individuals show evidence of extensive amyloid pathology yet little apparent clinical impairment, and others have a relatively low amyloid burden in the context of moderately advanced dementia. Transgenic rodent models utilising human familial Alzheimer’s disease gene mutations^3^ have been extensively used to model various aspects of APP/Aβ pathobiology but have not proved useful in exploring the mechanisms whereby this pathobiology affects disease pathogenesis and, as a consequence, we have no effective preclinical model of sporadic Alzheimer’s disease.

The advent of induced pluripotent stem cell (iPSC) technologies^4^ now makes it possible to derive patient-specific cell lines capable of differentiating into various cell types and thereby human cellular models of disease. Although familial Alzheimer’s disease iPSC-derived cells exhibit pathological APP-related phenotypes *in vitro*, sporadic Alzheimer’s disease iPSC-derived cells typically do not share the same phenotypes^5–7^. Recently however, iPSC-derived neurons were shown to display features *in vitro* that reflect analogous features from post-mortem material from the same individuals^8^. This has provided evidence on the feasibility of using individual cell models of disease to explore pathogenic mechanisms.

## Materials and methods

### Deep and Frequent Phenotyping (DFP) pilot cohort clinical data

The DFP pilot study protocol was previously published^9^, and a subset of the clinical outcomes, namely Mini Mental State Examination (MMSE) scores and global magnetoencephalography (MEG) readout, and participants was used for this iPSC study listed in Table 1. See Supplementary Materials for more details.

**Table 1:** Characteristics of the Deep and Frequent Phenotyping pilot cohort participants

| *Patients* | *MMSE score* | *APOE genotype* | Alzheimer's disease biomarkers |  | Clinical outcomes |  | Clinical vulnerability quotients<sup>§§§§</sup> |  |
| --- | --- | --- | --- | --- | --- | --- | --- | --- |
|  |  |  | *Cortical amyloid PET (SUVR)* | *CSF Aβ<sub>1-42</sub> concentration<sup>§</sup> (pg/ml)* | *MMSE score loss rate<sup>§§</sup> (per day)* | *Global MEG recording<sup>§§§</sup>* | *MMSE score loss rate quotients* | *MEG quotients* |
| 1 | 22 | $\epsilon 4/\epsilon 4$ | 1.60 | 298 | $1.5 \times 10^{-3}$ | 0.974 | 0.12 | 0.11 |
| 2 | 24 | $\epsilon 2/\epsilon 3$ | 1.21 | 269 | $8.4 \times 10^{-3}$ | 0.983 | 0.85 | 1.00 |
| 3 | 23 | $\epsilon 4/\epsilon 4$ | 1.28 | 344 | $6.3 \times 10^{-3}$ | 0.985 | 0.42 | 0.80 |
| 4 | 27 | $\epsilon 3/\epsilon 4$ | N/A | 395 | $0.2 \times 10^{-3}$ | N/A | 0.00 | N/A |
| 5 | 26 | $\epsilon 3/\epsilon 3$ | 1.63 | 254 | $0.8 \times 10^{-3}$ | 0.960 | 0.07 | 0.03 |
| 6 | 25 | $\epsilon 3/\epsilon 4$ | 1.60 | 450 | $3.9 \times 10^{-3}$ | 0.979 | 0.22 | 0.13 |
| 7 | 22 | $\epsilon 3/\epsilon 3$ | 1.28 | 252 | $3.7 \times 10^{-3}$ | 0.966 | 0.40 | 0.73 |
| 8 | 27 | $\epsilon 3/\epsilon 3$ | N/A | 236 | $1.2 \times 10^{-3}$ | N/A | 0.15 | N/A |
| 9 | 29 | $\epsilon 3/\epsilon 4$ | N/A | 262 | $3.0 \times 10^{-3}$ | N/A | 0.30 | N/A |
| 10 | 26 | $\epsilon 4/\epsilon 4$ | N/A | 164 | $2.9 \times 10^{-3}$ | N/A | 0.48 | N/A |
| 11 | 24 | $\epsilon 3/\epsilon 4$ | 1.30 | 414 | $1.9 \times 10^{-3}$ | N/A | 0.11 | N/A |
| 12 | 24 | $\epsilon 3/\epsilon 4$ | 1.62 | 287 | $10.5 \times 10^{-3}$ | 0.944 | 1.00 | 0.00 |
| 13 | 26 | $\epsilon 3/\epsilon 4$ | 1.56 | N/A | $3.1 \times 10^{-3}$ | N/A | N/A | N/A |
| 14 | 29 | $\epsilon 3/\epsilon 4$ | 1.18 | N/A | $0.5 \times 10^{-3}$ | 0.961 | N/A | 0.99 |

<sup>§</sup> Average value between two visits which were 169 days apart
<sup>§§</sup> Derived from MMSE score loss since estimated symptom onset and the baseline visit when the participants underwent MMSE
<sup>§§§</sup> Global efficiency metric from the γ-band (32-100 Hz)
<sup>§§§§</sup> Derived by dividing clinical outcome measurements with corresponding Alzheimer's disease biomarkers (i.e., MMSE score loss rate/CSF Aβ <sub>1-42</sub> and MEG/amyloid PET) and then rescaled to range from 0 to 1 within the DFP pilot cohort
Note: N/A – not available

### Patient iPSC-derived cortical neuronal culture

The patient iPSC lines were reprogrammed from peripheral blood mononuclear cells using Sendai virus. Subsequently, the iPSCs were differentiated into cortical neurons driven by *Ngn2* expression and plated in co-culture with primary rat astrocytes except for collecting conditioned media for the quantification of secreted Aβ (iPSC-derived neuronal monoculture). See Supplementary Materials for more details on the generation of iPSC, differentiation into cortical neurons and quantification of secreted Aβ.

### Synapse imaging

Immunocytochemistry was conducted on neurons treated with various extrinsic Aβ insults on Day 80 of the neuronal differentiation. Antibodies against presynaptic Synapsin I/II, postsynaptic HOMER1, and dendritic MAP2 were used. The samples were imaged on an Opera Phenix automated microscope, and the synapses were quantified relative to the total MAP2+ area to derive synaptic density for all downstream analyses. See Supplementary Materials for more details.

### Multi-electrode array (MEA) electrophysiology

The iPSCs were seeded directly onto the MEA plates for neuronal differentiation in co-culture with primary rat astrocytes. Baseline activities (2-min long) were recorded regularly from Day 45 onwards of the neuronal differentiation on a MEA equipment (Axion Biosystems, Maestro) with AxIS software v2.4.2.13 (Axion Biosystems). The neurons were treated with Aβ_1-42_ oligomers on Day 80 of the neuronal differentiation. See Supplementary Materials for more details.

### Statistical analyses

All statistical analyses were performed in GraphPad Prism 9.2.0. We reported Pearson’s coefficient of correlation and two-tailed *p*-values for correlations by simple linear regression. Kruskal-Wallis test was used for single-parameter comparisons amongst the patient lines. Welch’s *t*-test was used for the vulnerable-resilient group comparisons in the MEA experiment. * *p* < 0.05, ** *p* < 0.01, *** *p* < 0.001 and **** *p* < 0.0001.

### Data availability

Detailed raw data of the experiments are available from the corresponding authors upon reasonable request. The data from the DFP cohort can be requested via the Dementias Platform UK online portal (https://www.dementiasplatform.uk/research-hub/data-portal).

## Results

### iPSC lines from a comprehensively-phenotyped cohort of early Alzheimer’s disease patients

We set out to ask whether the heterogeneity of Alzheimer’s disease patients could be accurately reflected in iPSC models by comparing clinical outcomes *in vivo* with patient-derived neuronal phenotypes *in vitro*. We asked specifically whether clinical vulnerability to Aβ burden in the brain can be reflected by Aβ-induced cellular vulnerability in neurons derived from the same patients. In this study, we tapped into the comprehensive clinical datasets of the DFP pilot cohort^9^ (Table 1) from which we generated thirteen sporadic Alzheimer’s disease iPSC lines and one familial Alzheimer’s disease iPSC line (Patient #5) carrying an autosomal dominant *APP* mutation, to use in our experiments (Supplementary Table 1 and Supplementary Fig. 1). Previously, the DFP study has highlighted the heterogeneity of disease and also very considerable inter-individual variation in the impact of that amyloid pathology^10^. This suggests a difference in vulnerability or resilience in the face of amyloid pathology that might reflect differences either in the hypothesised amyloid cascade or in factors that interact with that cascade. Here, we seek to investigate if the functional consequences in response to Aβ burden in the brains of Alzheimer’s disease patients (instead of the accumulation of Aβ pathology *per se*^8^) can be recapitulated *in vitro* using iPSC models derived from the same patients.

### Levels of secreted Aβ_1-42_ from patient iPSC-derived neurons reflect the levels of donor CSF Aβ_1-42_

To understand if patient-derived iPSC models recapitulate the in-life clinical measures of their donors, we first differentiated all fourteen iPSC lines in parallel into cortical neurons in monoculture (Supplementary Fig. 2a) and showed that Aβ_1-42_ levels in the conditioned media correlate significantly and *negatively* with the same pathological Aβ species in the CSF from the patient donors (Fig. 1a), a characteristic phenomenon of Alzheimer’s disease patients thought to be due to the sequestration of Aβ_1-42_ in non-soluble cortical amyloid plaques^11^. Importantly, this relationship was not found for either Aβ_1-38_ or Aβ_1-40_ peptide comparisons and was not affected by the inclusion of the familial Alzheimer’s disease line. However, the Aβ_1-42_ / Aβ_1-40_, and Aβ_1-38_ / Aβ_1-40_ ratios were significantly increased in Patient #5 harbouring an *APP* mutation compared to the other patient lines, consistent with previous observation from another study^12^. This result provides further evidence that patient-derived neurons reflect the pathological features *in vivo* of that patient. We next went on to examine patient-specific cellular vulnerability to Aβ *in vitro*.

**Figure 1:**
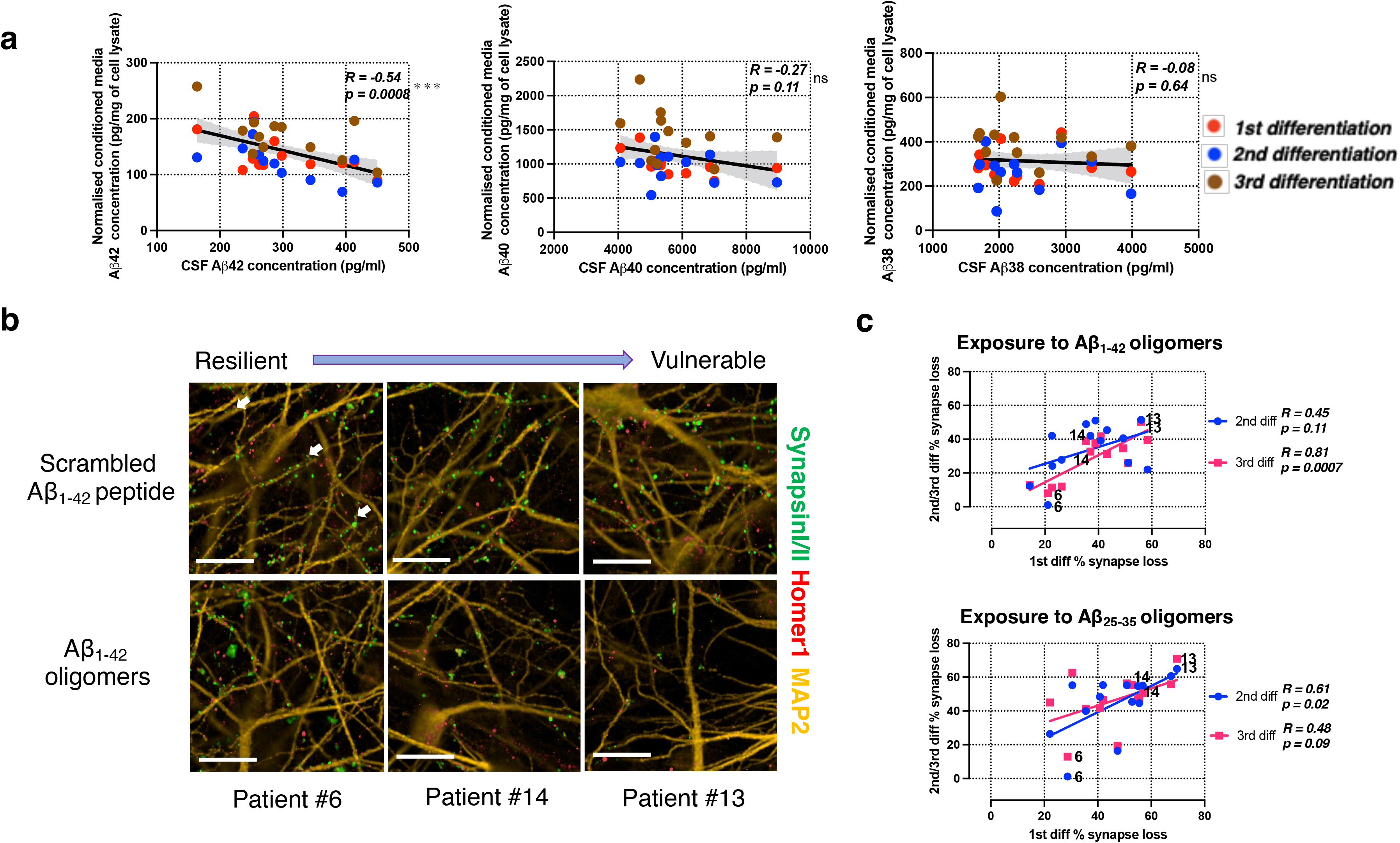
Levels of secreted Aβ from Alzheimer’s disease patient iPSC-derived cortical neurons correlated with patient CSF Aβ levels and extrinsic Aβ insults resulted in a spectrum of vulnerability of synapse loss in patient iPSC-derived cortical neurons. (A) Pairwise comparisons between the levels of secreted Aβ species from the patient-derived neurons and the levels of the same Aβ species in the patient CSF. Error band: 95% CI. *n* = 36 independent neuronal differentiation repeats per patient line. (B) Representative immunofluorescence images from three selected patient lines ranging from the least to the most vulnerable to Aβ_1-42_ oligomer insults relative to the scrambled peptide control treatment. The images are labelled with presynaptic (Synapsin I/II, green), postsynaptic (Homer1, red) and dendritic (MAP2, yellow) markers. White arrows indicate synapse examples with pre- and post-synaptic markers in apposition. Scale bar = 50 μm. (C) Pairwise comparisons of the degrees of synapse loss between neuronal differentiation repeats, caused by either Aβ_1-42_ or Aβ_25-35_ oligomers. The same three selected patient lines from Fig. 1b are highlighted in the graphs.

### Patient iPSC-derived neurons demonstrate a spectrum of synaptic vulnerability to Aβ insults

Dysregulation, and eventually loss, of synapses is one of the earliest pathological phenotypes of Alzheimer’s disease and leads to cognitive decline and memory loss^13, 14^. Electrophysiology, in particular MEG, is thought to be a surrogate of synaptic dysregulation and loss and hence provides an opportunity to explore whether the individual impact of Alzheimer’s disease pathology on synaptic health in people *in vivo* is reflected in their cells *in vitro*. We therefore sought to investigate synaptic vulnerability to Aβ insults *in vitro;* iPSC lines were again differentiated in parallel into cortical neurons, this time plated in co-culture with rat cortical astrocytes (Supplementary Fig. 2a, 2b). We then treated the neurons with a range of extrinsic Aβ insults listed in Table 2.

**Figure 2:**
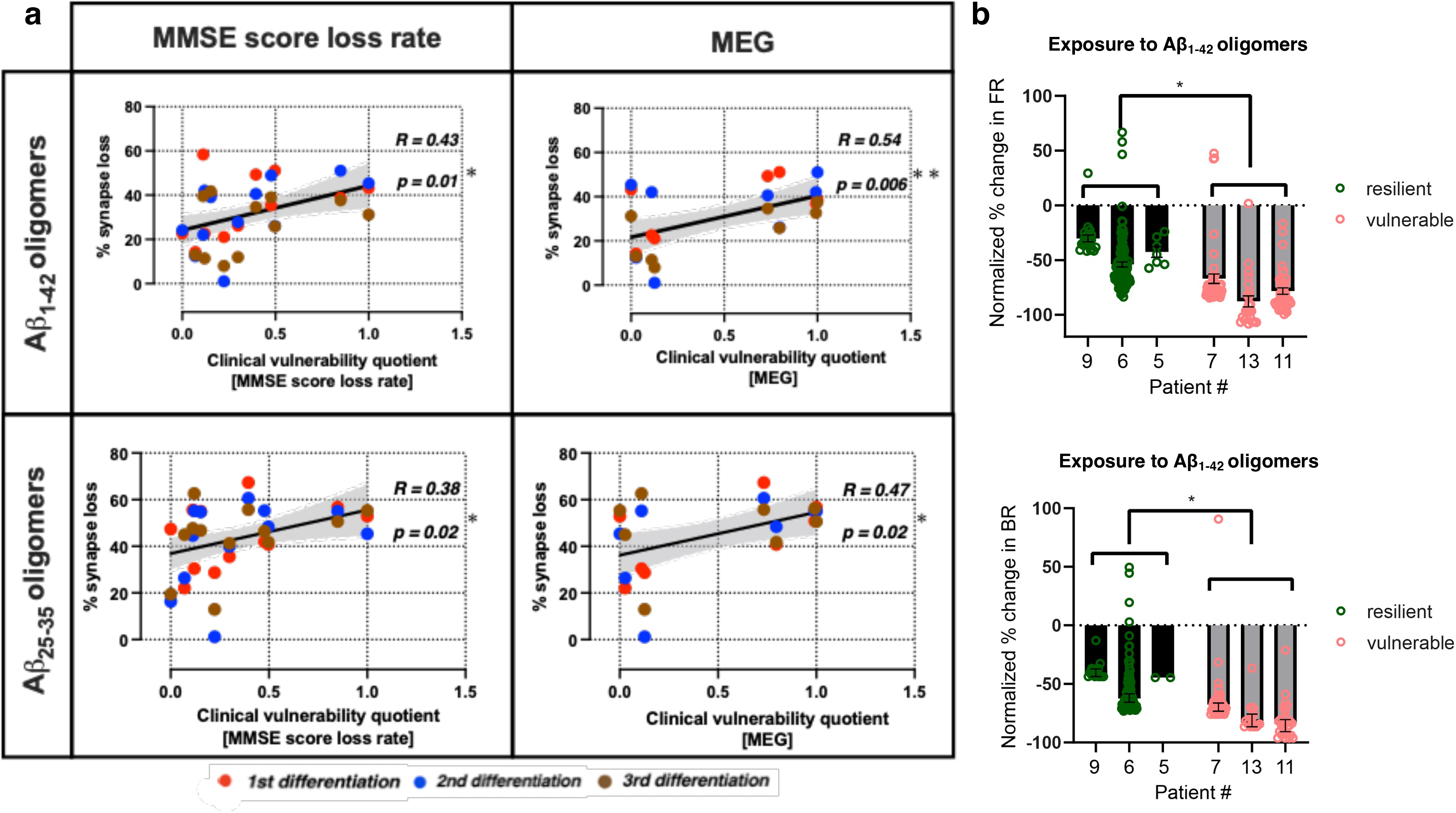
Synapse loss due to Aβ insults *in vitro* reflects clinical vulnerability in the same patients to Aβ burden *in vivo*. (A) Pairwise comparisons between the percentage of synapse loss and clinical vulnerability quotients. Each row denotes the type of oligomers used to induce synapse loss and each column denotes the selected clinical outcomes which have been corrected for Aβ_1-42_ concentration in the CSF (MMSE score loss rate) or amyloid PET SUVR (MEG). Error band: 95% CI. *n* = 35 (Aβ_1-42_ - MMSE score loss rate), 36 (Aβ_25-35_ - MMSE score loss rate) and 24 (MEG) independent neuronal differentiation repeats per patient line. (B) Comparison of the resilient group (Patients #9, #6 and #5; green) and vulnerable group (Patients #7, #13 and #11; red) neuronal response (Day 80 of the neuronal differentiation) in their firing rate (FR) and burst rate (BR) to Aβ_1-42_ 10 μM after two days of incubation. The vulnerable group showed greater decrease in activity compared to the resilient group in both FR and BR. Each datapoint represents an electrode recording. *n* = 22 (#9), 114 (#6), 7 (#5), 49 (#7), 24 (#13), and 41 (#11) for the FR data whereas *n* = 17 (#9), 93 (#6), 2 (#5), 43 (#7), 18 (#13), and 29 (#11) for the BR data. Percentage change from baseline was normalised against changes of untreated media control. Mean ± SEM; Welch’s *t*-test was used for statistical analysis.

**Table 2:** Types of extrinsic Aβ insults used to cause synapse loss in iPSC-derived cortical neurons

| Type of Aβ insult | Dosage | Aβ treatment control | Pathological relevance |
| --- | --- | --- | --- |
| Aβ<sub>1-42</sub> oligomers | 10 μM for 24 h | Scrambled Aβ<sub>1-42</sub> peptides | Elevated levels in Alzheimer's disease brain; major pathological Aβ species found in amyloid plaques<sup>1,2</sup> |
| Aβ<sub>25-35</sub> oligomers | 20 μM for 24 h | Aβ<sub>35-25</sub> (reversed) peptides | Represents the biologically active region of Aβ; shortest fragment that exhibits large β-sheet aggregated structures and retains the toxicity of the full-length peptide<sup>3</sup> |
| Alzheimer's disease brain homogenate | 25% v/v for 72 h | Artificial CSF | Derived from post-mortem brain tissues; composition of Aβ species that most closely recapitulates the pathological milieu |

All three exogenous Aβ treatments resulted in decreased synaptic density in all patient-derived cortical neurons relative to control treatments. However, the different patient lines showed different levels of impact of Aβ insults on synapse loss, allowing us to rank lines from the most resilient to the most vulnerable (Fig. 1b and Supplementary Fig. 2c, 3). Notably, cellular vulnerability in the patient carrying the familial Alzheimer’s disease *APP* mutation that generated the most endogenous Aβ_1-42_ was within the range, but was relatively resilient to the impact of exogenous Aβ insults. All neurons displayed functional activity by firing action potentials on Day 80 of neuronal differentiation (Supplementary Fig. 2d). The synapse loss datasets demonstrated good reproducibility over three repeat independent iPSC differentiations. By comparing the extent of synapse loss between differentiation repeats, we confirmed that the specific levels of vulnerability in each line of iPSC-derived neurons in response to Aβ insults remained consistent across all differentiation repeats (Fig. 1c and Supplementary Fig. 4a, 4c). Importantly, similar patient line-specific vulnerability measured by synapse loss was also consistent across the different Aβ insults used, especially between Aβ_1-42_ and Aβ_25-35_ oligomers where there is a significant and positive correlation (Supplementary Fig. 4b). A positive correlation was also observed across differentiation repeats when the neurons were treated with Alzheimer’s disease brain homogenate (Supplementary Fig. 4a, 4c). The synapse loss data indicated that the degree of synapse loss due to the exposure to extrinsic Aβ in functional cortical neurons is patient-specific, cell-autonomous, and reproducible across insults and differentiation repeats.

### Synaptic vulnerability to Aβ insults in vitro reflects clinical vulnerability to Aβ burden in vivo

Next, we explored if the levels of synaptic vulnerability to Aβ insults in the patient-derived neurons *in vitro* was a reflection of the individual’s response to amyloid in life as measured using electrophysiological measures of synaptic activity and measures of cognitive decline, the ultimate clinical manifestation of synaptic dysfunction. While in the *in vitro* experiments the cells were exposed to the same amount of Aβ insult, *in vivo* the individuals showed a range of amyloid burden. Global MEG recordings and cognitive decline measured by MMSE score loss rate (Table 1) were therefore adjusted as a function of the patients’ individual levels of Aβ burden measured by amyloid PET standard uptake value ratios (SUVR) and CSF Aβ_1-42_ concentration, respectively. This yielded a personal ‘clinical vulnerability quotient’ representing the synaptic and cognitive response as a function of amyloid pathological load per individual. The resultant quotients were then rescaled within the DFP pilot cohort to range from 0 (least vulnerable or most resilient) to 1 (most vulnerable, least resilient) thereby facilitating comparisons between the MMSE loss rate clinical vulnerability quotients and the MEG clinical vulnerability quotients.

Using this analysis, we found that the amount of synapse loss in patient-derived neurons caused by Aβ insults *in vitro* reflects the personal clinical vulnerability to Aβ burden *in vivo,* whether measured by the surrogate measure of synaptic number and function, MEG, or by cognitive decline, the core clinical manifestation of synaptic loss. Specifically, we observed a positive correlation between the percentage of synapse loss caused by both Aβ_1-42_ and Aβ_25-35_ oligomers and clinical vulnerability quotients, demonstrating that greater cellular vulnerability correlates significantly with greater clinical vulnerability in these patients (Fig. 2a). Synapse loss due to the exposure to human Alzheimer’s disease brain homogenate resulted in similar correlation with clinical vulnerability quotients (Supplementary Fig. 5).

We then selected the three most vulnerable together with the three most resilient patient lines and investigated whether their electrophysiological activities were also differentially affected based on their synaptic vulnerability *in vitro*. As for the synaptic loss measures, the neurons derived from the most vulnerable patient lines exhibited greater reductions of firing and burst rates caused by the exposure to Aβ_1-42_ oligomers as compared to the most resilient patient lines (Fig. 2b). The scrambled Aβ_1-42_ peptide control did not elicit any change in the levels of neuronal activities (Supplementary Fig. 6). Additionally, the differences in synapse loss in the patient-derived neurons could not be explained by their *APOE* variants (Supplementary Fig. 7) nor by the single case of an *APP* mutation carrier who scored as both relatively resilient to amyloid *in vivo* and to Aβ *in vitro* suggesting that the resilience/vulnerability to Aβ is not driven either by the most significant genetic variant associated with sporadic Alzheimer’s disease or by mutations in the *APP* gene itself.

In conclusion, we show here that neurons derived from Alzheimer’s disease patients retain the same individual vulnerability to Aβ that the person from whom they were derived, showed using both biomarkers and clinical measures that reflect the synaptic phenotypes measured *in vitro*.

## Discussion

In this study, we demonstrate for the first time that cellular vulnerability to Aβ insults *in vitro* reflects clinical vulnerability to Aβ burden *in vivo,* specifically in people living with Alzheimer’s disease dementia, by establishing the correlation between synapse loss in individual Alzheimer’s disease patient-derived neurons and their clinical outcomes. This was further supported by neurons from the more vulnerable group of patients exhibiting more deleterious response to extrinsic Aβ insults as measured by their levels of neuronal activity as compared to the resilient group. This approach of integrating clinical in-life data with disease modelling in the laboratory presents a tractable method of Alzheimer’s disease modelling with iPSCs.

Decline in cognition estimated from time since onset and current cognitive score, and ‘brain activity’ assessed using MEG were selected as clinical outcomes likely to be reflections of synaptic health and so broadly analogous to the synaptic loss data we measured *in vitro*. We report here that the amount of cognitive decline as a function of amyloid burden correlates with more severe Aβ-driven synapse loss and loss of synaptic function, as measured using MEA electrophysiology, in the patient-derived neurons. Although it has been known that synapse loss correlates with cognitive decline in Alzheimer’s disease^14, 15^, and that MEG identifies neurophysiological changes that are specific to Alzheimer’s disease, it remains unclear how different brain MEG signals change at different stages of Alzheimer’s disease progression^16, 17^. Interestingly, we find a clear correlation between *greater* brain activity levels measured by MEG correlating with more severe Aβ-driven synapse loss in the patient-derived neurons. This apparently counterintuitive observation is in line with a considerable amount of evidence for hyperexcitability in the early phases of Alzheimer’s disease. Neurons exhibit hyperactivation particularly during the mild cognitive impairment stage before hypoactivation as disease progresses^18, 19^, and hyperexcitability leading to seizure activity is increased in Alzheimer’s disease, perhaps as a function of amyloid related pathology^20^. Our findings substantiate the role of hyperexcitability in early Alzheimer’s disease and provide a model with which to explore such therapeutics discovery.

It has recently been shown that several measures of secreted Aβ peptides in iPSC-derived cortical neurons from Alzheimer’s disease patients reflect the extent of Aβ neuropathology of their donors^8^. We extend that previous work on post-mortem, end of life, neuropathological findings to in-life, early in disease, clinical measurements by showing that the levels of Aβ_1-42_ secreted from patient-derived neurons correlate with the levels of the same pathological Aβ species in the patient CSF samples (Fig. 1a). However, we have now shown that not only is there a correlation between cellular phenotypes and analogous phenotypes in post-mortem brain and in patients, but that the functional consequences of those phenotypes – the response to Aβ as well as the amount of Aβ – are preserved in the cells. Crucially, the inclusion of the familial Alzheimer’s disease case did not affect the cellular-clinical correlation in vulnerability to Aβ. The neurons from the familial Alzheimer’s disease individual in fact belong to one of the more resilient patient lines *in vitro* even though this individual has the greatest Aβ burden measured by amyloid PET within this cohort, further reinforcing our interpretation that the iPSC models specifically reflect the vulnerability to Aβ measured by clinical outcomes instead of the levels of Aβ accumulation in the brain.

In conclusion, we reveal that cellular vulnerability reflects clinical vulnerability to Aβ in Alzheimer’s disease by modelling with patient iPSC-derived neurons and integrating cellular and clinical data from a highly-phenotyped cohort. We first demonstrated the correlation between levels of Aβ_1-42_ secreted from patient iPSC-derived cortical neurons and the levels of the same pathological Aβ species in the patient CSF samples, and then demonstrated Aβ-driven synapse loss and dysfunction in iPSC-derived neurons reflects relevant clinical outcomes as a function of Aβ burden in the brain. This work establishes the feasibility of modelling in-life Alzheimer’s disease clinical phenotypes with patient iPSC-derived neurons. Beyond that, as we can model inter-individual variability in clinical response to Aβ insult in an individual’s own iPSC derived neurons *in vitro*, this raises the potential for interrogating mechanisms, and identifying targets for precision therapy in human cell models.

## Abbreviations

Aβ: Amyloid-β

BR: burst rate

DFP: Deep and Frequent Phenotyping

FR: firing rate

iPSC: induced pluripotent stem cells

MEA: multi-electrode array

MEG: magnetoencephalography

MMSE: Mini Mental State Examination

SUVR: standard uptake value ratio.

## Acknowledgements

We thank all participants of the Deep and Frequent Phenotyping pilot cohort for contributing clinical data and samples towards this study. We thank S Cowley, J Vowles and S Chintawar for technical assistance in characterising iPSC lines.

## Author contributions

R.W.-M., Z.M.C. and S.L. conceived the study design. B.N., S.L. and R.W.-M. contributed to the experiment design. S.L. was the PI and D.C, T.C., D.f., R.N.G., E.K., J.L., P.M., B.H.R., J.B.R, A.J.T. and G.Z. contributed to the design and execution of the clinical study. B.N., H.A.R. and T.W. contributed materials for experiments, performed experiments, data acquisition, analysis, and presentation. K.A., E.M.H. and A.H. contributed to cell line generation, quality control and expansion. B.N., H.A.R., T.W., I.K., E.M.R., N.J.B., J.R., Z.M.C., S.L. and R.W.-M. contributed to data analysis and interpretation. E.M.R., N.J.B., Z.M.C, S.L. and R.W.-M. supervised the study. B.N. drafted the manuscript and B.N., T.W., I.K., N.J.B., Z.M.C., S.L. and R.W.-M. edited the manuscript. R.W.-M. finalised the manuscript before all authors approved the final version of the manuscript.

## Ethics statement

The DFP cohort study was approved by the London Central Research Ethics Committee, 14/LO/1467. The human iPSC lines used for this study were derived from human blood erythroblasts, (NHS Research Ethics Committee: 10/H0505/71) and were derived as part of the Innovative Medicine Initiative-EU sponsored StemBANCC consortium. Informed consent was obtained from all donors.

## Supplementary Materials

**Supplementary Table 1:** DFP pilot cohort patient-derived iPSC IDs

| **Supplementary Table 1: DFP pilot cohort patient-derived iPSC IDs** |  |
| --- | --- |
| **Patient #** | **iPSC ID** |
| 1 | BPC-927 03-01 |
| 2 | BPC-928 03-07 |
| 3 | BPC-929 03-07 |
| 4 | BPC-932 03-03 |
| 5 | BPC-933 03-12 |
| 6 | BPC-934 03-02 |
| 7 | BPC-936 03-07 |
| 8 | BPC-937 03-01 |
| 9 | BPC-939 03-01 |
| 10 | BPC-940 03-08 |
| 11 | BPC-943 03-03 |
| 12 | BPC-944 03-04-01A |
| 13 | BPC-946 04-10 |
| 14 | BPC-947 04-09 |

---

**Supplementary Figure 1:**
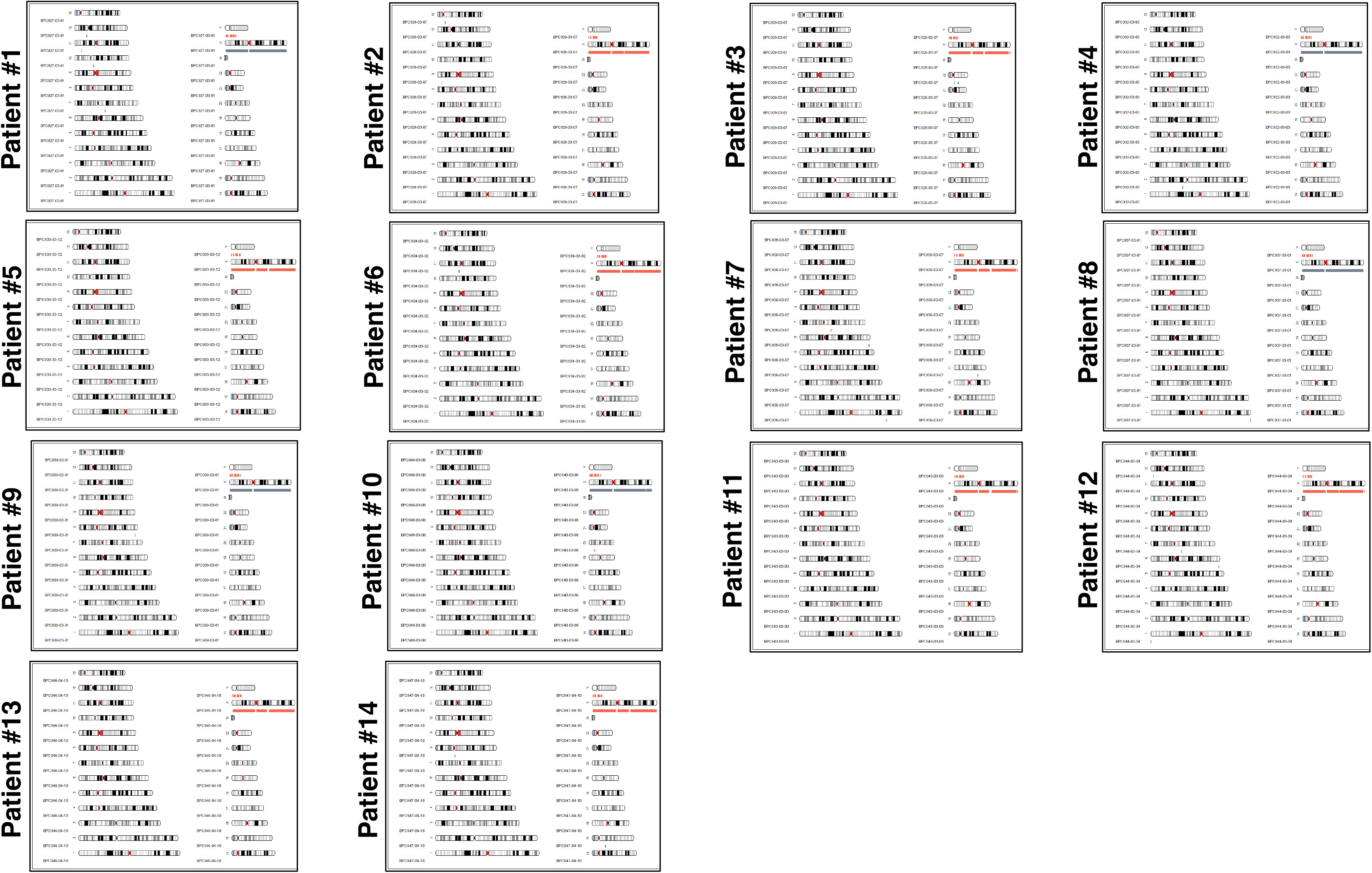
Alzheimer’s disease patient-derived iPSC quality controls. Genome integrity of the Alzheimer’s disease patient-derived iPSC lines, examined by the Illumina OmniExpress24 single nucleotide polymorphism array. Karyograms (KaryoStudio, Illumina) show amplifications (green)/deletions (orange)/loss of heterozygosity regions (grey) alongside the relevant chromosome. Female X chromosome is annotated in grey.

**Supplementary Figure 2:**
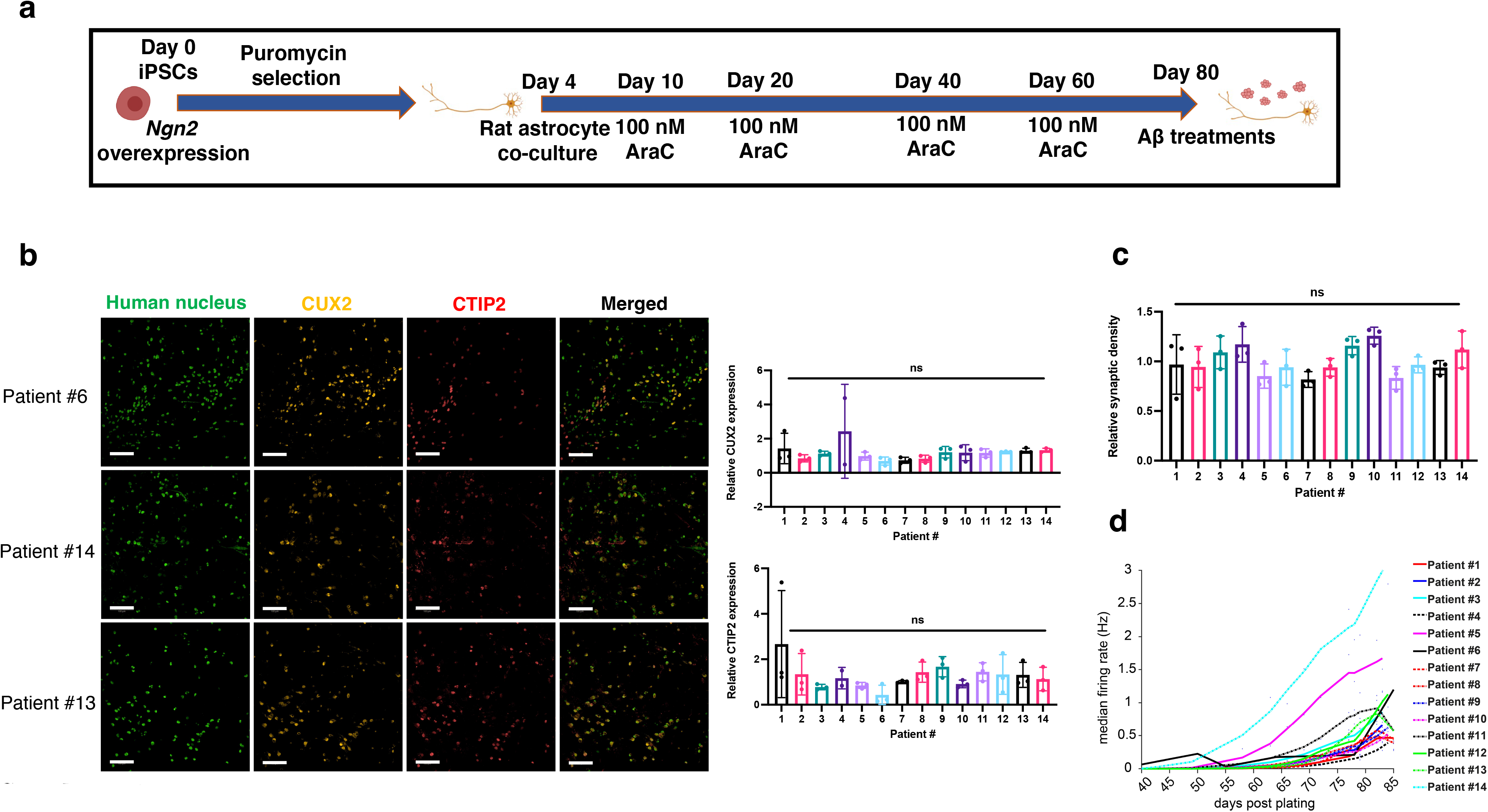
Characterisation of iPSC-derived cortical neurons. (A) Schematic of the cortical neuron differentiation protocol over 80 days. (B) Representative images of cortical markers from three patient lines ranging from the least to the most vulnerable to Aβ insults, as well as the quantification of relative expression levels across all patient lines. Scale bar = 100 μm. Mean ± SD. *n* = 3 independent neuronal differentiation repeats. (C) Relative synaptic density across all patient-derived cortical neurons, normalised to the mean of synaptic density per neuronal differentiation. Mean ± SD. *n* = 3 independent neuronal differentiation repeats. (D) Neuronal activity increase over time measured by MEA from one neuronal differentiation. The figure plots smoothed line (the lowest function in MATLAB) of extracellular firing rate medians in Hz of cortical neurons between Day 40 to Day 85 post plating on the MEA plate. The small dots are the raw data points recorded. Each raw recording has the length of 2 min from which median was calculated.

**Supplementary Figure 3:**
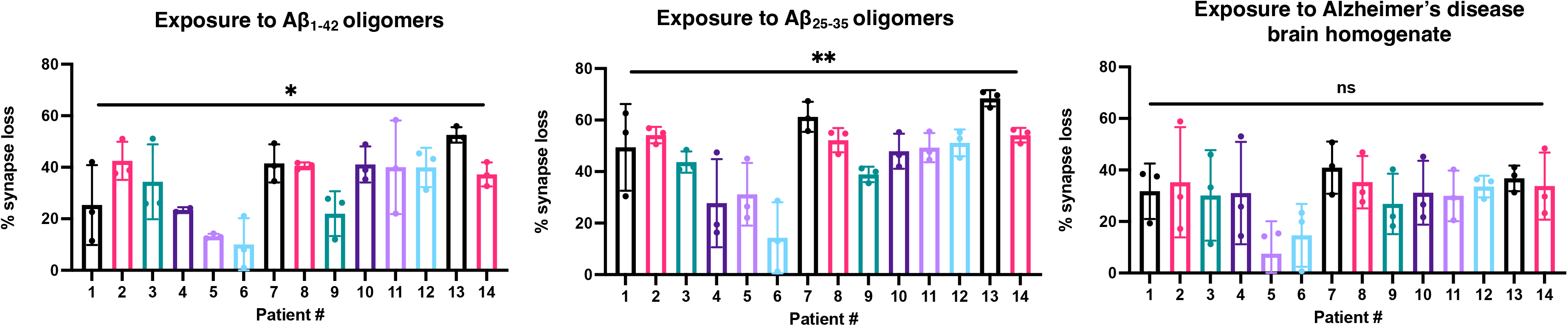
Varying levels of synapse loss caused by Aβ insults. Percentage of synapse loss caused by Aβ_1-42_ oligomers, Aβ_25-35_ oligomers and Alzheimer’s disease brain homogenate across all patient lines normalised to their respective treatment controls. Mean ± SD. *n* = 3 independent neuronal differentiation repeats.

**Supplementary Figure 4:**
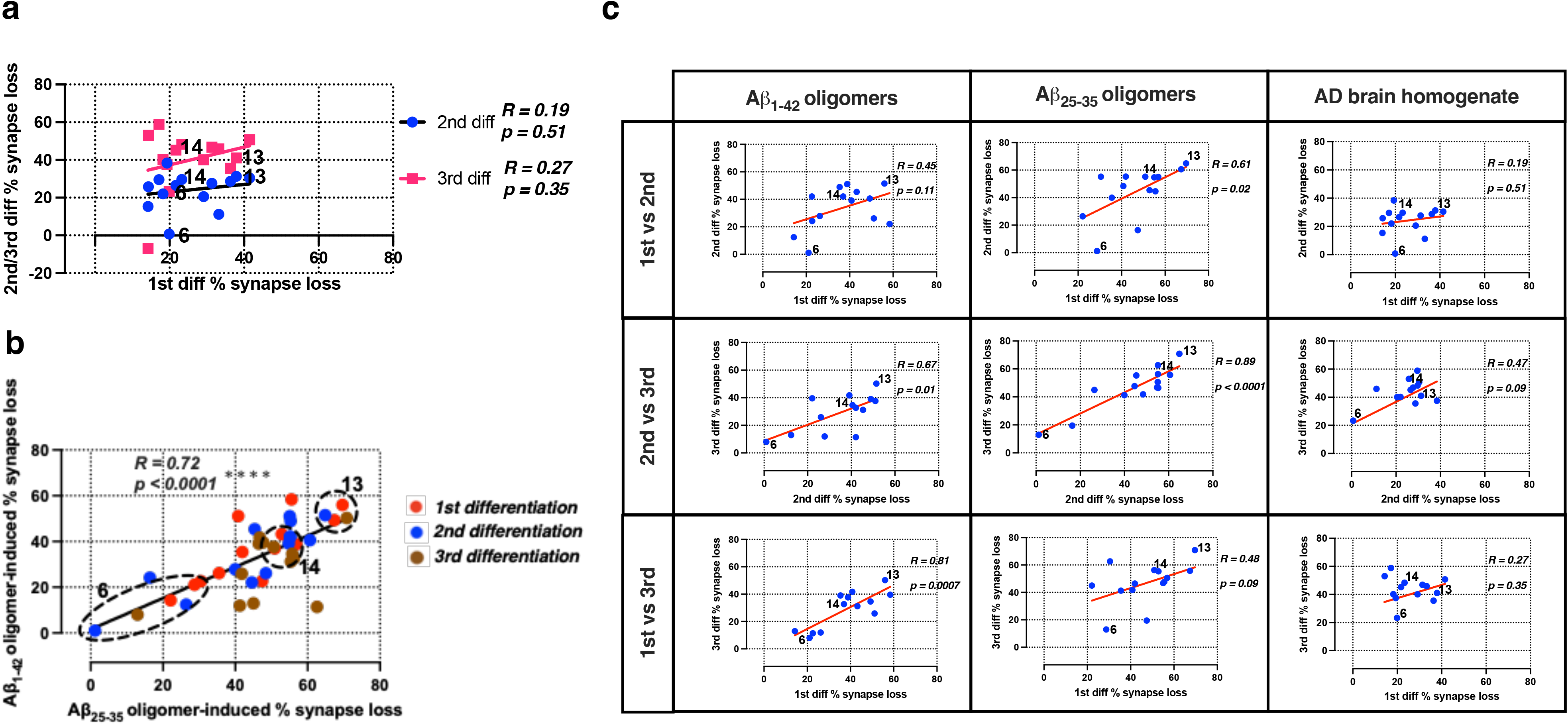
Good reproducibility of the synapse loss data across neuronal differentiation repeats indicates cell-autonomous vulnerability to Aβ insults. (A) Pairwise comparisons on the degrees of synapse loss between differentiation repeats caused by Alzheimer’s disease brain homogenate. The same three selected patient lines from Fig. 1b are highlighted in the graphs. (B) Pairwise comparison on the degrees of synapse loss caused by either Aβ_1-42_ or Aβ_25-35_ oligomers. The zones where the same three selected patient lines from Fig. 1b can be found are circled in the graphs. (C) Breakdown of individual pairwise comparisons on the degrees of synapse loss between differentiation repeats summarised in Fig 1c. Each row denotes the two differentiation repeats in question and each column denotes the Aβ insult used to induce synapse loss.

**Supplementary Figure 5:**
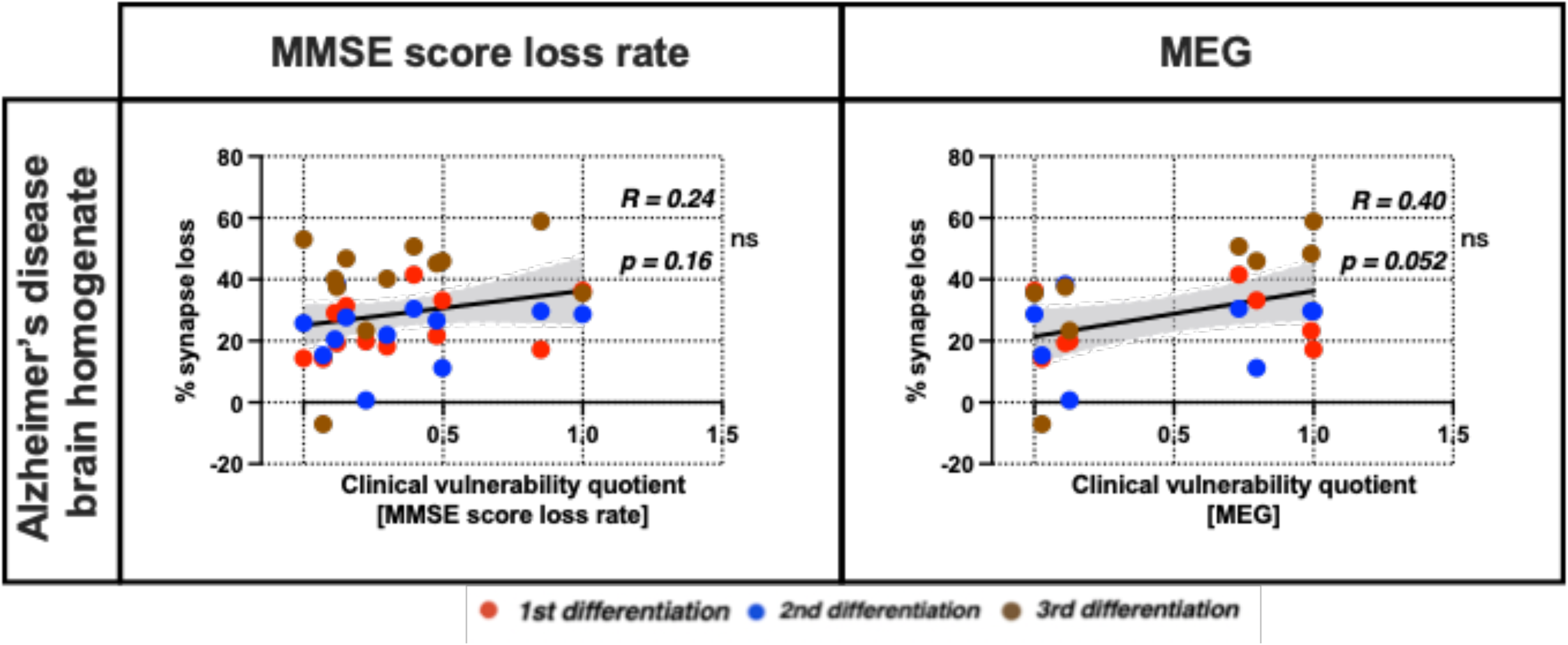
Correlations between the synapse loss data caused by Alzheimer’s disease brain homogenate and clinical vulnerability quotients. Pairwise correlations between the percentage of synapse loss caused by Alzheimer’s disease brain homogenate *in vitro* and clinical vulnerability quotients on MEG (normalised to amyloid PET SUVR) and MMSE (normalised to the concentration of Aβ_1-42_) outcomes *in vivo*. Each column denotes the selected clinical outcomes. Error band: 95% CI. *n* = 36 (MMSE score loss rate/CSF Aβ_1-42_) and 24 (MEG/PET) independent neuronal differentiation repeats per patient line.

**Supplementary Figure 6:**
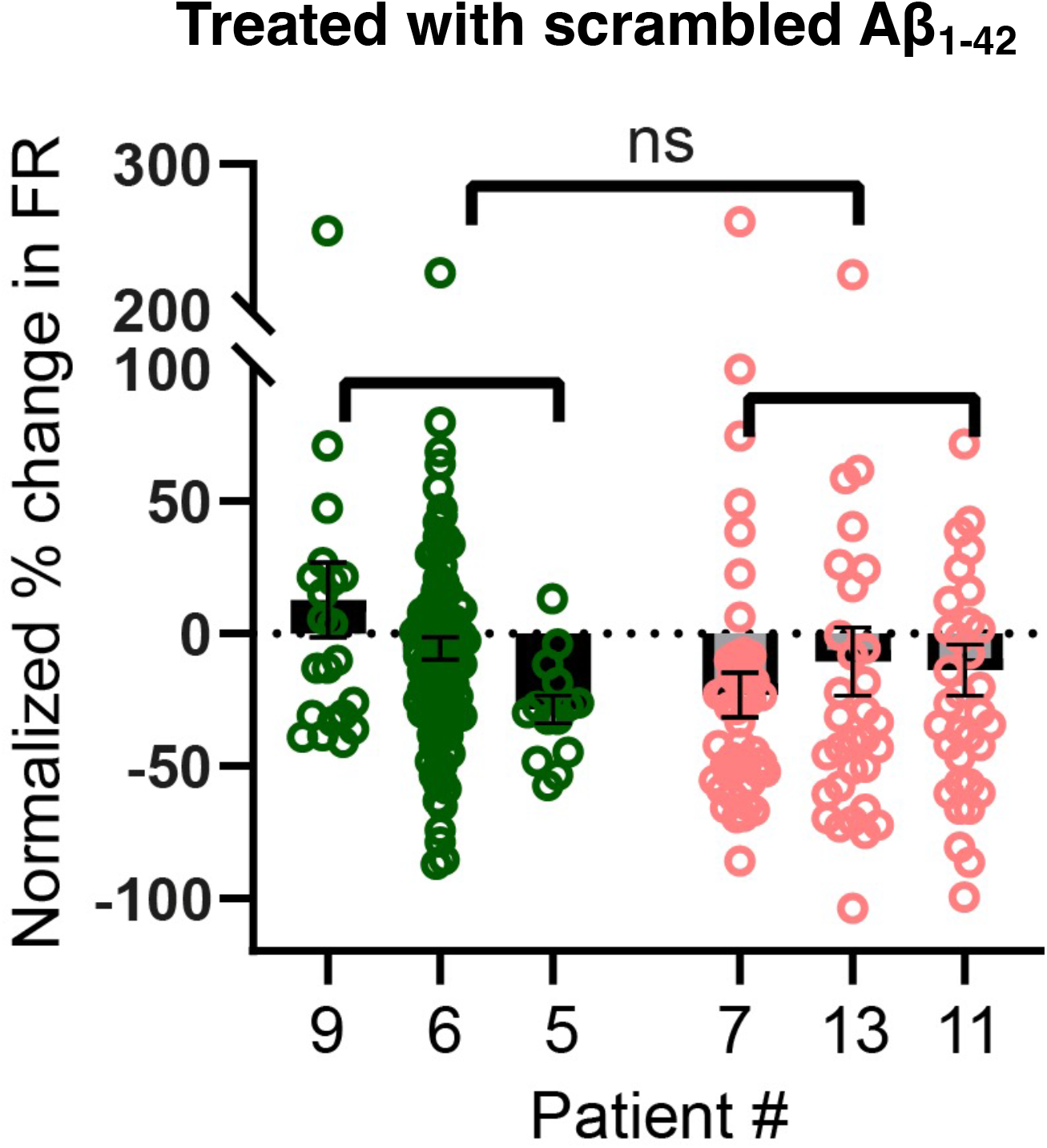
Scrambled Aβ_1-42_ treatment did not cause any electrophysiological changes measured by MEA. Comparison of the resilient group (Patients #9, #6 and #5; green) and vulnerable group (Patients #7, #13 and #11; red) neuronal response in their firing rate (FR) to scrambled Aβ_1-42_ 10 μM on the second day of incubation. Each datapoint represents an electrode recording. *n* = 22 (#9), 104 (#6), 14 (#5), 46 (#7), 33 (#13), and 35 (#11). Percentage change from baseline was normalised against changes of untreated media control. Mean ± SEM; Welch’s *t*-test was used for statistical analysis.

**Supplementary Figure 7:**
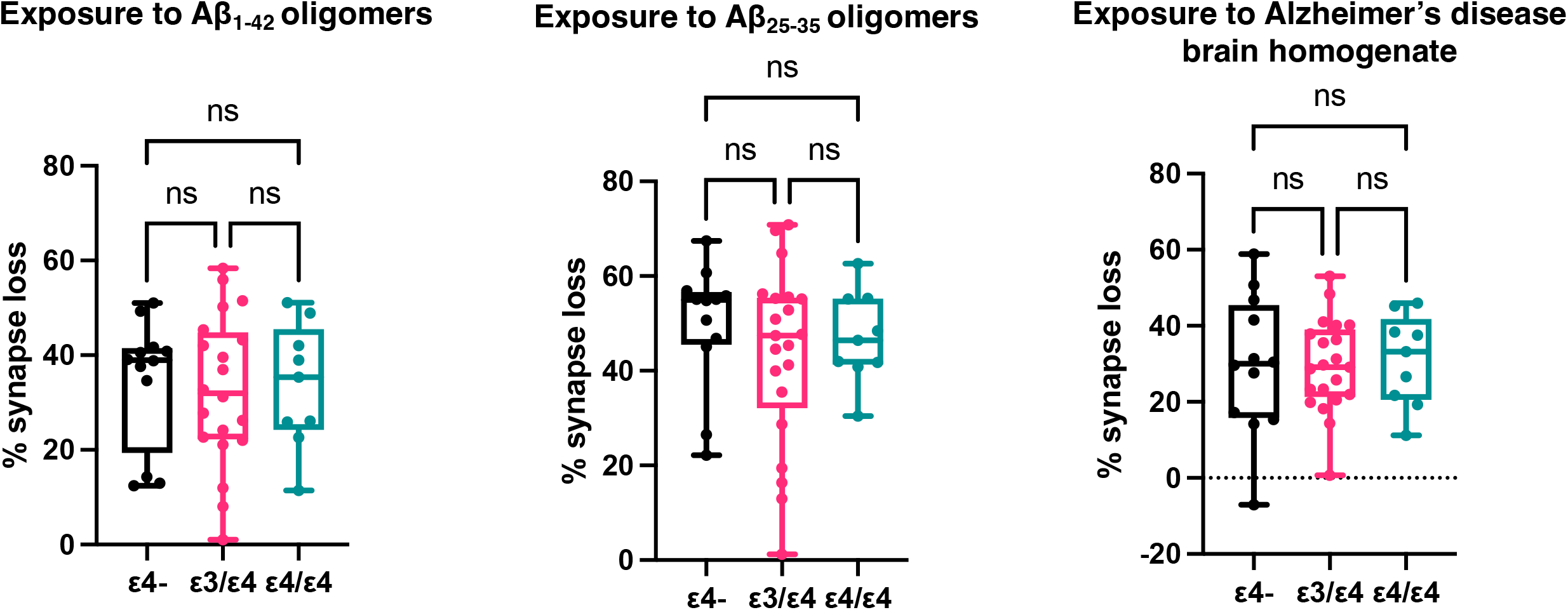
*APOE* genotypes could not differentiate the synaptic vulnerability to extrinsic Aβ insults *in vitro*. Box plots (centre line, median; box limits, interquartile range; whiskers, data range; points, all data points) showing the percentage of synapse loss caused by Aβ_1-42_ oligomers, Aβ_25-35_ oligomers and Alzheimer’s disease brain homogenate with patients distinguished by their *APOE* variant genotypes. *n* = 12 (ε4-), 20 (ε3/ε4) and 9 (ε4/ε4) independent neuronal differentiation repeats per patient line.

## Supplementary materials and methods

All reagents were purchased from Sigma unless stated otherwise. All iPSC-derived neuronal culture incubated at 37°C and 5% CO_2_.

### Deep and Frequent Phenotyping cohort pilot study and clinical data

The Deep and Frequent Phenotyping (DFP) cohort pilot study protocol was previously published^1^ and a subset of the comprehensive clinical data and study participants (fourteen early symptomatic Alzheimer’s disease cases) was used for the analyses. These participants were recruited based on their clinical assessment meeting the criteria of early Alzheimer’s disease. The participant ages were in the age range groups of 51-60 yeas (2/14), 61-70 years (4/14), 71-80 years (6/14), and 81-90 years (2/14) and 5/14 were female. One participant with a family history of familial Alzheimer’s disease was a known carrier of an *APP* mutation, the remainder had a family history compatible with sporadic Alzheimer’s disease. Briefly for the study protocol, both amyloid PET imaging with [^18^F] AV45 (0–60 min, 150 ± 24 MBq) and magnetoencephalography (MEG) recordings were conducted once in 10/14 and 8/14 of the pilot study participants, respectively. Global efficiency metric from the γ-band (32-100 Hz) of the MEG raw data was used for analysis as it is the oscillation range linked to cognitive function and local connectivity^2^. Lumbar puncture was performed over two visits 169 days apart in 12/14 of the study participants for CSF collection and Aβ_1-42_ peptide concentration was quantified via electrochemiluminescence in 96-well plates from Meso Scale Discovery (Aβ peptide panel 1 with 6E10 antibody), before deriving the average values from the two visits for downstream analyses. All pilot study participants underwent cognitive testing including a Mini Mental State Examination (MMSE; Mean = 25.3) and a MMSE score loss rate measuring cognitive decline was derived by dividing the MMSE score loss since estimated symptom onset and baseline visit to the clinic with the time since estimated symptom onset in days.

### Generation of Alzheimer’s disease patient-derived iPSC lines from blood samples

Blood samples (8 ml) were remixed via gently inversion and centrifuged at 1800 g/20 min with brakeless deceleration. The plasma portion was removed, taking care not to disturb the whitish phase ring containing the peripheral blood mononuclear cells (PBMC). PBMC were diluted to 40 ml using phosphate-buffered saline (PBS; Thermo) (added dropwise to prevent osmotic shock) and centrifuged at 300 g/15 min. Cells were counted and plated at 5 × 10^6^/ml in Expansion I medium which consists of StemSpan SFEM (Stem Cell Technologies) supplemented with lipid concentrate (1%. Gibco), dexamethasone (1 µM), IGF-1 (40 ng/ml, R&D Systems), IL-3 (10 ng/ml, R&D Systems), EPO (2 U/ml, R&D Systems) and SCF (100 ng/ml R&D Systems). The remaining wells were filled with PBS to maintain a humid atmosphere (continued throughout all expansion and reprogramming steps). From DIV-1 to DIV-6, a 50% media change (Expansion I medium) was performed. Erythroblasts should appear ∼ DIV-5.

To purify the erythroblast population, 4 ml Percoll was first added to a 15 ml tube. The wells were washed with DMEM (used for all washing steps) before a maximum of 8 ml cell solution was slowly trickled onto the Percoll solution to collect erythroblasts. The solution was centrifuged at 1000 g/20 min with brakeless deceleration. The supernatant above the phase ring was transferred to a new tube and centrifuged at 300 g/5 min and washed twice to remove the Percoll. Purified erythroblasts were plated at 1-1.5 × 10^6^/ml in Expansion II medium which has the same constituents as Expansion I medium except IL-3. On DIV-8/9, erythroblasts were collected, centrifuged at 200 g/5 min, resuspended in Expansion II medium, and plated at 1-1.5 × 10^6^/ml to prevent cells differentiating down the erythroid lineage.

Before reprogramming erythroblasts to iPSCs, each well of a six-well plate was coated with 1 ml of 0.1% gelatine at 37°C for > 20 minutes. Mitomycin-C treated CF1 Mouse Embryonic Fibroblasts (MEF) were thawed and transferred to a tube containing 34 ml MEF medium which consists of Advanced DMEM supplemented with fetal calf seum (10%), GlutaMAX (1%) and 2-mercaptoethanol (0.1%) all purchased from Life Technologies. The gelatine was aspirated from the wells and two ml of MEF suspension were added per well. Plates were incubated overnight at 37°C before erythroblasts were plated after undergoing viral transduction.

Erythroblasts were collected and centrifuged at 200 g/5 min when they were ready to be infected with Sendai viruses expressing Yamanaka factors. The pellet was resuspended in Expansion II media. 1.5 × 10^5^ cells were collected and made up to 200 μl in Expansion II media. An aliquot from the CytoTune™-iPS 2.0 Sendai Reprogramming Kit (Thermo) was thawed on ice, mixed with 150 μl Expansion II media and added to cell suspension. The entire suspension was transferred to a well in a 24-well plate. Viral supernatant was removed 23 h later by collecting cells and centrifuging at 300 g/4 min. The pellet was resuspended in Expansion II media and transferred to a well in a 24-well plate before incubating for 48 h.

Finally, MEF medium was removed from feeder plates which were washed with PBS before 1 ml of Expansion II media was added. The transduced erythroblasts were collected, centrifuged at 300 g/4 minutes, and resuspended in Expansion II media. The cells were plated at a range of densities (1.5 - 4.5 × 10^4^/ml) which yielded approximately eight to twelve clones but allowed the clones to grow large enough for picking without overcrowding. A 50% media change was performed on the following days with the following media – DIV-5 (Expansion II media), DIV- 7/8 (hES medium which consists of KnockOut DMEM supplemented with 20% KnockOut Serum Replacement, 1% GlutaMAX, 1% non-essential amino acids, 0.1% 2-mercaptoethanol and 10 ng/ml BFGF) and DIV-10 (Conditioned Medium derived from MEF culture with hES medium). Clones appeared ∼ DIV-15 and were picked ∼ DIV-22. If clones did not appear by DIV-40, the line was deemed to have failed to reprogramme. Colonies that displayed embryonic stem cell-like morphology were selected via manual picking. All iPSC lines used in this study express pluripotency markers Tra-1-60 and NANOG measured by fluorescence-activated cell sorting.

### Maintaining iPSC culture and differentiation into iPSC-derived cortical neurons

iPSC culture was maintained by growing the cells on Matrigel matrix (Corning) and feeding with mTeSR^TM^ medium (STEMCELL technologies) which was replaced daily. We differentiated the iPSC lines into cortical neurons by overexpressing Neurogenin-2 (*Ngn2*)^3^. All 14 iPSC lines were infected with lentivirus carrying the plasmids for *Ngn2* overexpression, before we plated the cells onto poly-ornithine (100 µg/ml) plus laminin (10 µg/ml) coated cell culture plates at 60,000 cells/cm^2^ (double for several lines which did not grow well) in mTeSR^TM^ medium (STEMCELL Technologies) supplemented with Y-27632 (Tocris) at 10 µM on Day 0. The mTeSR^TM^ medium was replaced with Neurobasal^TM^ medium (Gibco) supplemented with B27^TM^ (Thermo), GlutaMAX^TM^ (Gibco), Penicillin-Streptomycin (Gibco), neurotrophin-3 (10 ng/ml). BDNF (10 ng/ml, Peprotech), doxycycline (1 µg/ml), laminin (200 ng/ml) and ascorbic acid (200 µM) five hours after plating. Subsequently, the cell culture medium was further supplemented with puromycin (1.5 µg/ml) on Day 2 only.

The cells underwent the only and final passage on Day 4 with Accutase^TM^ and were plated at 25,000 cells/cm^2^ onto a confluent layer of rat cortical astrocytes (Thermo Fisher) in half-area 96-well plates. The rat cortical astrocytes were introduced to facilitate neuronal maturation^4,5^, improve neuronal morphology for imaging assays, and improve cell attachment to withstand subsequent biochemical procedures. Half-feeding took place twice per week from Day 4 onwards with the abovementioned B27-containing medium. Finally, we also supplemented the medium with Ara-C (100 nM final concentration) on Day 10, 20, 40 and 60.

### Multi-electrode array (MEA)

The iPSCs were seeded directly onto the MEA plates, and 30,000 rat cortical astrocytes were seeded into each well of the MEA plates on Day 5 of the differentiation. From Day 45 onwards of the cortical neuron differentiation, 2-min long recordings were taken after 5 mins plate settling time on the MEA reader regularly over time (Axion Biosystems, Maestro) with AxIS software v2.4.2.13 (Axion Biosystems). The plate was kept at 37°C while recordings were taken. The raw recording files were then extracted with AxIS software (Axion Biosystems) and processed with custom script in MATLAB (version R2020b). Firing rate (FR) is defined as number of extracellular electrical spikes in ms window per recording length above noise (> 6 STD). Burst rate (BR) is defined as number of groups of minimum 5 spikes with ISI < 100ms counted per recording length.

### Meso Scale Discovery immunoassay of Aβ peptides

The iPSC derived neurons were grown as described previously without the Day 4 passage onto rat astrocytes until Day 40. Cell conditioned media was collected after 48 hours and stored at −80°C. Cells were washed once with PBS, and M-PER™ (Thermo) added for 20 min on ice. Cell suspension was centrifuged at 14,000 g for 10 min at 4 °C. The supernatant was collected, and protein concentration was quantified by BCA assay (Thermo). Measurement of Aβ38, Aβ40, Aβ42 was measured by electrochemiluminescence using Meso Scale Discovery V-PLEX Aβ peptide panel (6E10), which was carried out according to manufacturer’s protocol. Cell media samples were run in triplicate, with 25 µg of each cell lysate run in duplicate and kept on a plate shaker covered with a plate seal at room temperature during incubation. The Meso Scale Discovery Workbench 4.0 software was used to analyse Aβ levels. Conditioned media samples were normalised to the average of total protein concentration in the lysate.

### Oligomerisation of Aβ peptides and treatment in neuronal culture

Both lyophilised Aβ_1-42_ and treatment control scrambled Aβ_1-42_ peptides (Bachem, H-1368 and H-7406) were resuspended to 1 mM in Hexafluoro-2-propanol. The tubes were vortexed and left sitting at room temperature for 30 min, before they were aliquoted and dried in a Speed-Vac concentrator for 30 min. We kept the Aβ film at −80°C. To oligomerise the Aβ_1-42_ peptides, we first resuspended the Aβ film in DMSO to 5 mM before sonicating in water bath for 10 min. PBS was then added to result in 100 µM solution and the tubes were left stationary at 4°C for 24 h. Just before treating the cells with Aβ oligomers, the solution was centrifuged at 14,000 g for 10 min at 4°C to remove any precipitate/fibrils.

Both lyophilised Aβ_25-35_ and treatment control Aβ_35-25_ peptides (Bachem, H-1192 and H-2964) were resuspended to 2 mg/ml in deionised water and vortexed before incubating at 37°C for 2 h for oligomerisation. The vial was then aliquoted and frozen at −80°C.

The iPSC-derived neurons were incubated with Aβ oligomers for 24 h before paraformaldehyde fixation.

### Human Alzheimer’s disease brain homogenate extraction

The extraction protocol of human Alzheimer’s disease brain homogenate was modified from a published method^6^. We sourced the post-mortem frozen frontal cortices from two Alzheimer’s disease patients (Patient #1: 73yo, female, Braak stage VI, 75 h post-mortem delay; Patient #2: 81yo, male, Braak stage VI, 26 h post-mortem delay) from the Oxford Brain Bank. We first thawed the brain tissues on ice prior to homogenisation with Dounce homogenisers for 25 strokes in cold artificial CSF (aCSF: 124 mM NaCl, 2.8 mM KCl, 1.25 mM NaH_2_PO_4_ and 26 mM NaHCO_3_, pH = 7.4) with a ratio of 1 g of tissue to 4 ml of aCSF supplemented with a panel of protease inhibitors (5 mM EDTA, 1 mM EGTA, 5 ug/ml leupeptin, 5 µg/ml aprotinin. 2 µg/ml pepstatin, 120 µg/ml Pefabloc and 5 mM NaF). The homogenisation was followed by centrifugation at 200,000 g for 110 min at 4°C and the supernatant was transferred into a Slide-A-Lyzer™ G2 Dialysis Cassettes 2K MWCO in 100 times volume of aCSF without protease inhibitors for 72 h. The aCSF was replaced every 24 h and the resultant aliquots were frozen at −80°C.

The iPSC-derived neurons were incubated with either 25% Alzheimer’s disease brain homogenate (1:1 mixture from the two cortices) or aCSF without protease inhibitors as the treatment control in the cell culture medium (v/v) for 72 h at 37°C before paraformaldehyde fixation.

### Immunocytochemistry

Adherent neurons were fixed in 4% paraformaldehyde for 5 min, followed by treating with 0.5% saponin in PBS for 20 min for permeabilisation. To block the samples, we treated the plates with 10% normal goat serum with 0.01% tween-20 in PBS for 30 min. Primary antibodies were then left incubating with the samples at 4°C overnight with 1% normal goat serum and 0.01% tween-20, before washing with PBS 3 times. Secondary antibodies were then applied in 1% normal goat serum and 0.01% tween-20 at room temperature for 1 h before washing for another four times. The primary antibodies we used were: Guinea pig anti-SYNAPSIN I/II (Synaptic Systems, 1:500), rabbit anti-HOMER1 (Synaptic Systems, 1:500), chicken anti-MAP2 (Abcam, 1:1000), mouse anti-human nuclear antigen (Abcam, 1:200), rabbit anti-CUX2 (Abcam, 1:200) and rat anti-CTIP2 (Abcam, 1:500). The secondary antibodies we used were: Goat anti-guinea pig Dylight 488 (Abcam), goat anti-mouse Alexa Fluor 488, goat anti-rabbit Alexa Fluor 555, goat anti-chicken Alexa Fluor 555, goat anti-rabbit Alexa Fluor 647 and goat anti-rat Alexa Fluor 647 (Thermo) at 1:1000 dilution.

### High-content imaging and analysis

#### Synapse

The 96-well plates were imaged on the Perkin Elmer Opera Phenix high-content imager. We captured 15 images per well with a 43X objective at +1 µm focus level with the binning value of 1. We then analysed the image with the Harmony software v4.9 from Perkin Elmer with a customised pipeline. The MAP2-positive neurites were identified with 0.5 overall threshold as the region of interest and resized by expanding outwards by 5 px to cover synaptic signals which lay slightly above the MAP2 signals. Both presynaptic (SYNAPSIN I/II) and postsynaptic (HOMER1) signals were then identified with Method A of the “Find spots” function with threshold values of 0.17 and 0.14, respectively. We also filtered away the spots which were larger than 100 px^2^. Finally, the synapses were ascertained by finding HOMER1 signals in the vicinity of SYNAPSIN I/II signal regions which had been resized by expanding outwards by 5 px. The absolute number of synapses was then normalised to the total MAP2-positive area to derive synaptic density which was used for all downstream analyses. All the values of synaptic density downregulation due to the Aβ extrinsic insults were then normalised to the corresponding treatment controls i.e., Aβ_1-42_ normalised to scrambled Aβ_1-42_, Aβ_25-35_ normalised to Aβ_35-25_ (reversed), and Alzheimer’s disease brain homogenate normalised to aCSF.

#### Cortical markers

We captured 15 images at −1, 0 and +1 and µm focus levels per well with a 20X objective and binning value of 2. We analysed the images on the same Harmony software by first identifying human nuclei among the co-culture with rat astrocytes and filtering away the nuclei with circularity less than 0.6. The percentage of cortical marker-positive cells was calculated by selecting the human nuclei with cortical marker mean signal intensity greater than a threshold which was determined as the mean intensity across all human nuclei. Finally, we derived relative cortical marker expression by normalising the percentage of cortical marker-positive neurons to the geometric mean across all fourteen patient lines.

